# From Few Labels to Full Insight: 3D Semantic Segmentation of Brain White Matter Microstructures from a Sparsely Annotated X-ray Nano Holotomography Dataset

**DOI:** 10.1101/2025.06.08.658301

**Authors:** Mahsa Amirrashedi, Hans Martin Kjer, Miren Lur Barquin Torre, Tim B. Dyrby

## Abstract

Morphological analysis of white matter (WM) microstructures provides invaluable insights into its function and brain health overall. Recently, synchrotron X-ray Nano Holography (XNH) imaging has gained momentum in microstructural studies owing to its unique combination of capabilities, including a wide field of view (FOV), nanoscale resolution, and fast volumetric acquisition. However, the 3D segmentation of tiny and densely packed microstructures within the WM, the very first step toward quantification, presents a significant bottleneck. Herein, we address this challenge using an expert-in-the-loop 3D deep learning (DL) framework. The initial model was a 3D U-Net architecture trained on sparsely annotated data, exclusively from the corpus callosum (CC), a region characterized by relatively simple axonal morphology and parallel orientation of axonal bundles. The high-confidence predictions of the model (p > 0.7) from both CC and the complex crossing fiber regions are then refined by an expert and reintegrated into the training process to expand the training dataset and the corresponding labels. Building upon this foundation, we benchmarked alternative architectures, including 3D convolutional neural networks (CNN) and transformer-based architectures, alongside different loss functions. We showcased the segmentation performance of the models in XNH volumes with varying microstructural morphology and organization. All DL architectures achieved satisfactory performance with an overall segmentation accuracy > 0.96 and an average Dice score > 0.8 in 3D segmentation of axons, blood vessels, cells, and vacuoles. This study presents a multi-class 3D segmentation of WM microstructures in large FOVs up to ∼202 × 202 × 780 µm^3^ scanned through XNH technology. Our framework not only accelerates high-fidelity segmentation in challenging datasets but also paves the way for fast quantitative analysis in large-scale 3D neuroimaging studies.

## 1. Introduction

Morphological characterization of white matter (WM) microstructures holds the key to understanding its functional dynamics both in normal and disease conditions. Gaining such insights demands imaging techniques capable of delivering nanometer (nm)-scale resolution across millimeter (mm)-scale field of view (FOV). Moreover, since anatomy and its microstructures (axons, cells, blood vessels) are in 3D, so should the imaging modalities be. Currently available imaging modalities, however, pose significant challenges in achieving such insights. For instance, Magnetic Resonance Imaging (MRI) and Light Microscopy (LM) lack the nanoscopic image resolution required for in-depth microstructural analysis. While high-resolution imaging methods, such as Electron Microscopy (EM), can achieve resolutions ranging from 0.5 Å to a few nm [1], they are limited by small FOVs (a few µm) and painstaking data collection processes. These include serial sectioning and subsequent slice alignment to reconstruct 3D volumes [2]. Recently, synchrotron X-ray Nano Holotomography (XNH) imaging of intact samples has gained momentum in large-scale microstructural studies and is a fast-developing imaging technology [2, 3, 4, 5]. Today, beamlines like ID16A at European Synchrotron Research Facility (ESRF), offer XNH for a wide FOV (hundreds of µm) with isotropic image resolution toward 40 nm and facilitate volumetric acquisition of mm-scale specimens within just few hours and further volume extension via stitching, without further need for invasive tissue preparation such as slicing, staining, and clearing [2]. The larger FOV of XNH could be used to obtain a basic understanding of anatomy’s spatial organisation in 3D [5] and validate biophysical modelling of microstructure in MRI studies [6] with potential translation to clinical applications. This unique combination of capabilities positions XNH as a powerful imaging modality for studying complex brain microstructures in their native 3D context, thereby allowing for volumetric visualization and investigation of intact, large-scale features such as long-range axons, cellular arrangement within the axonal pathways, and the vascular network [4]. Nonetheless, quantitative assessment of WM micromorphology necessitates instance-level segmentation of individual microstructures within. Manual annotation of these tiny and densely packed microstructures is highly demanding and time-consuming. This often leads to sparse and incomplete annotations of XNH images [4]; hence, a realistic, full statistical description of all anatomical features cannot be obtained and will only be limited to what has been manually or semi-automatically annotated [4]. Traditional image processing algorithms for segmentation also fall short in handling the complexity and variability of 3D microstructural features across different brain regions, where tissue organization and microstructural morphology change [4, 5]. All together, these challenges underscore the pressing need for automated solutions.

Recent studies have demonstrated the potential of deep neural networks in the automatic segmentation of WM microstructures, with most focusing on axons, the dominant microstructure in WM. For instance, the AxonDeep tool was developed initially for 2D segmentation of axons [7]. The author explored supervised and semi-supervised techniques using fully connected networks (FCN) and generative adversarial networks (GAN). The methods applied to 2D LM images of murine optic nerves yielded the best results via the GAN framework, trained semi-supervised, with a Dice score of 0.81 [7]. Another example is AimSeg [8], a supervised method implemented in Ilastik [9] and designed to segment three different WM microstructures, such as axons, myelin, and inner tongue of the myelin wrapping in 2D. Using AimSeg, F1-scores of 0.71, 0.85, and 0.83 were achieved for axons, myelin, and inner tongue on 2D EM images acquired from the mouse corpus callosum (CC), respectively [8]. Another notable framework is AxonDeepSeg, which utilizes a U-Net to segment axons in 2D microscopy data [10]. The method was evaluated on EM images from different species, and Dice scores of 0.9 and 0.94 were reported for axonal segmentation in macaque and mouse CC, respectively [10]. Some studies have also examined the utility of 2D U-Net for segmenting unmyelinated peripheral axons in EM data [11]. Critically, axons are inherently 3D anatomical structures with morphologies varying along their length [4]. Moreover, 2D imaging techniques can not sufficiently represent 3D objects. Toward this end, few efforts have sought to extend segmentation into the 3D domain. The DeepACSON framework [12] was developed to segment 3D EM datasets in the mouse CC and cingulum, where two separate networks were trained: one for axons/mitochondria segmentation and another for cell nuclei segmentation. Using DeepACSON, F1-scores of 0.886 and 0.861 were reported for myelin and myelinated axons, respectively [12]. More recently, a U-Net-based method was employed to segment short association axons in a 3D multibeam serial EM dataset acquired from the human temporal lobe. The author trained several sets of 2D and 3D U-Nets for various tasks, including but not limited to segmentation and artifact detection, and achieved Dice scores of ∼0.95 in axon and myelin segmentation [13].

While previous studies have provided valuable insights into axonal segmentation, none have demonstrated 3D multi-class segmentation of diverse microstructures within the WM. Moreover, previous works have only relied on a single WM region without assessing the performance of their DL model on different pathways of WM, where both morphology and microstructural organization change. Herein, we aimed to address these gaps by introducing an expert-in-the-loop 3D DL framework capable of segmenting different microstructures in the WM, including axons, cells, vacuoles, and blood vessels, from a large FOV of 3D XNH images ∼202 × 202 × 780 µm^3^, similar to an MRI voxel. Unlike prior studies focusing on a single WM region with relatively simple axonal arrangements, such as the CC, our dataset also includes a deep WM region with complex axonal morphology and organization, where up to three axonal populations intersect [5]. For initialization, we only use sparsely annotated datasets available from a region of straight axons, including a few axons obtained from [4]. By incorporating an expert-in-the-loop strategy, we demonstrate extending the anatomical feature space to segment a broader range of morphological sizes within the resolution limits of the dataset.

In the second phase of the study, after expanding the labeled dataset by incorporating the expert-in-the-loop strategy, we sought to train the most performant DL model in our segmentation context. To do so, we trained and benchmarked alternative DL architectures, including state-of-the-art 3D convolutional neural networks (CNNs) and transformer-based architectures, along with three different loss functions. We specifically evaluate the importance of the patch size (input to the network) in securing robust segmentation and demonstrate how it limits the maximal 3D geometrical features that can be accurately segmented; hence, the importance of balancing the patch size with the sizes of the features to be segmented.

## 2. Materials and Methods

### 2.1. Dataset

Datasets used in this study are publicly available in [4, 5]. Detailed information regarding sample preparation and XNH scanning is described in [4]; however, a brief summary is provided here for clarity and readability. Samples used in this study were derived from a 32-month-old female vervet monkey brain. The monkey was housed and cared for on the island of St. Kitts in compliance with a protocol authorized by the Caribbean Primate Center of St. Kitts [4]. The monkey brain was perfusion-fixed and sourced from the Montreal Monkey Brain Bank. The entire monkey brain was sectioned into sagittal slices with a thickness of 2-4 mm. Samples from the splenium region of the CC and the centrum semiovale (CS), among others, were extracted with a 1 mm diameter biopsy punch and fixed with 2.5% glutaraldehyde. These samples were stained with 0.5% osmium tetroxide (OsO_4_), dehydrated using an alcohol series, and embedded in Epon resin. These Epon-embedded samples were then trimmed to 1 × 1 × 4 mm^3^ blocks and scanned at beamline ID16A of ESRF via a nano-focused X-ray beam of 17 keV. Angular holographic projections were collected at four distinct propagation distances, with sample rotation over 180°. These projections were aligned to reconstruct the phase maps of the samples. Each tomographic scan included 1800 projections and was recorded with an exposure time of 0.22 seconds. In total, each scan required approximately 4 hours.

Samples from two WM regions with different microstructural characterization were included in this study: 1) the splenium of CC, which is composed of straight axons running parallel to each other and connecting the brain hemispheres [4], and 2) CS, a deep WM region including up to three crossing fiber populations [5]. Opposed to CC, CS consists of more tortuous and complex axonal morphology, with axons running in different directions. XNH scans for each of the WM regions mentioned above consisted of 4 consecutive volumes (n = 4 for CC and n = 4 for CS), referred to here as volumes #1 to #4. Each of these volumes was reconstructed with an isotropic voxel resolution of 75 nm for CC and 100 nm for CS with an image matrix of 2048 × 2048 × 2048. A relatively high level of noise was observed in the XNH images, due in part to issues related to tissue preparation (e.g., scratches on the sample, which affected the phase-contrast signal). We followed the same processing method described in [4] to reduce the image noise. Each volume was subjected to Gaussian filtering and downsampled by a factor of 5, resulting in dimensions of 410 × 410 × 410. After downsampling, the final resolution was 375 nm for CC and 500 nm for CS. To further extend the FOV, the downsampled volumes from each region were stitched together (Fig. 1A). The resulting extended volumes cover 153.75 × 153.75 × 574.5 µm^3^ for CC and 202 × 202 × 780 µm^3^ for CS.

**Figure 1:**
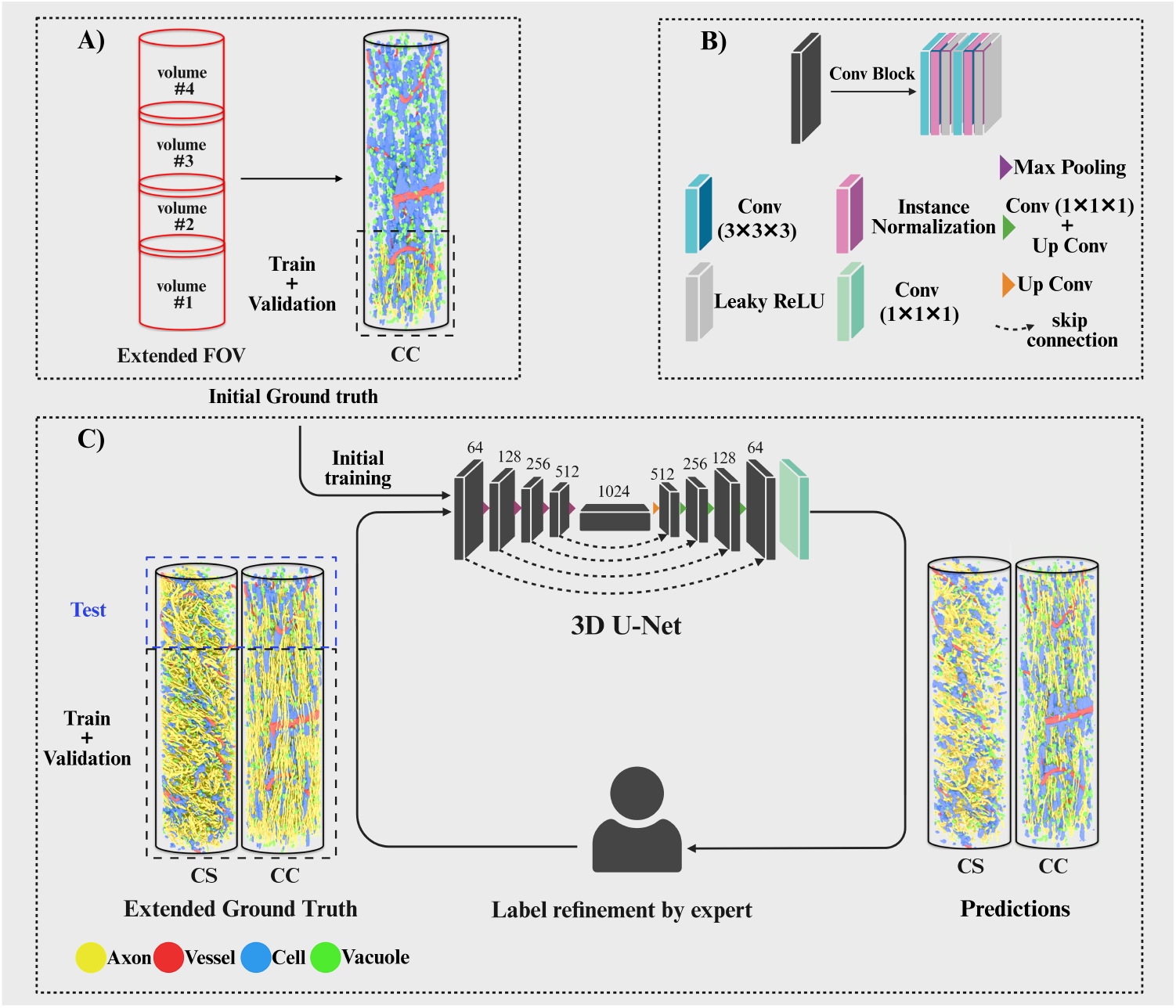
The expert-in-the-loop deep learning workflow for semantic segmentation of various microstructures in the white matter. (A) Extended XNH volume from the corpus callosum (CC) and the initial ground truth (GT). The extended field of view (FOV) was achieved by stitching four consecutive XNH volumes with a small overlap. The initial GT labels for the axons were limited to 54 axons annotated manually in volume #1 of CC, while annotations for the other microstructures were available from the extended CC volume. These include annotations for whole blood vessels, the majority of cells, and vacuoles. Annotations for cells include both single cells and cell clusters. (B) Convolutional blocks of the 3D U-Net model. These consist of (3 × 3 × 3) convolutions, instance normalization, Leaky ReLU activations, max pooling, and up-convolutions. (C) The expert-in-the-loop workflow. The 3D U-Net is initially trained using the sparse GT labels from volume #1 of the CC. Predictions were performed on the extended CC and centrum semiovale (CS) volumes. These were then refined by the expert, resulting in an extended GT(EGT) used for the next rounds of training and evaluation. Volume #4 from each region was held out as an unseen test and used only for post-hoc assessment of the model performance. This figure was created in BioRender under the agreement number PW28CPY3E8.

### 2.2. Ground truth labeling

For the first round of training (hereinafter referred to as initial training), we utilized the sparse annotations freely available in [4]. These 3D annotations were limited to the CC region and served as ground truth (GT) labels. As illustrated in Fig. 1A, entire blood vessels, the majority of vacuoles, and cells in the extended CC volume were annotated. Note that GT labels for cells included both single cells and cell clusters of varying sizes. However, only a few annotations were available for the axons, which were limited to one specific volume of the CC (volume #1 of CC). These manually drawn axonal labels [4], totaling 54 axons, were biased towards larger splenial axons with a mean diameter of ∼2.5 µm and a maximum length of 170 µm.

### 2.3. Extended GT labels and the expert-in-the-loop strategy

Due to the sparseness of the GT labels (particularly the axons), we adopted an expert-in-the-loop training strategy to extend the GT labels, here referred to as extended ground truth (EGT). As shown in Fig. 1C, the expert-in-the-loop training follows two main steps. First, a 3D U-Net model was trained on 3D patches sampled randomly from volume #1 of CC, where GT labels for all target classes were available (*n*_volume_ = 1). In subsequent training rounds, all predicted labels with high confidence scores (*p* > 0.7) from extended CC and CS volumes (Fig. 1C) were verified and manually corrected by the expert (M.A.). The ITK-SNAP software (V 4.0.1) [14] was used for visual inspection and label correction. Additional adjustments were also made for false positives (FP) and false negatives (FN) using morphological operations and volume thresholding in Python 3.10.10. Note that in each training round, the refined labels were added to the EGT from the previous round, and the model was trained from scratch using the updated EGT. By doing so, we aimed to reduce the risk of catastrophic forgetting. Finally, we expanded the training and validation sets by sixfold (*n*_volume_ = 6) and covered a broader range of anatomical feature sizes without requiring manual 3D annotation from scratch. In all training rounds, a 70/30 split was used for training and validation. To evaluate the performance of the model, a single volume (#4) from each region (CC and CS) was held out as a test dataset. The test dataset was not included in the expert-in-the-loop procedure and was only used for post-hoc analysis.

It is worth noting that the tissue sample was not homogeneously stained throughout [4, 5]; some regions exhibited a weak tissue intensity contrast compared to fully stained areas. This uneven staining of the samples made dense segmentation of cells, vacuoles, and axons impractical. Therefore, we focused our evaluation exclusively on well-stained microstructures that the expert reliably delineated and corrected.

### 2.4. Model architectures

We studied three different convolutional neural networks (CNNs), namely modified U-Net [15], Attention U-Net [16], SegResNetVAE [17], and also a transformer-based architecture, Swin UNETR[18]. The implementations of all models were performed in 3D using PyTorch [19] and MONAI [20] libraries.

### 2.5. U-Net

We used a 4-stage modified 3D U-Net architecture with a bottleneck. The model features a symmetrical encoder-decoder structure and skip connections [15]. The encoder consists of sequential convolutional layers, where each layer employs two (3 × 3 × 3) convolutional blocks followed by instance normalization and Leaky Rectified Linear Unit (Leaky ReLU) activation, and dropout regularization. In the encoder, downsampling is performed via max pooling. The decoder mirrors the encoder, where upsampling is achieved via transposed convolution and the concatenation of features from the corresponding encoder layers. The initial number of filters is set to 64, which doubles at each downsampling stage and reaches 1024 in the bottleneck layer. To enable multi-scale learning and calculate auxiliary losses at different decoder levels, the model produces outputs at four different decoding stages via (1 × 1 × 1) convolutional layers.

#### 2.4.2. Attention U-Net

Attention U-Net [16] has the same architecture as U-Net, but with additional attention gates implemented before skip connections [16]. These attention gates refine the encoder features and modulate them based on relevance to the decoder’s upsampled guidance signal. Dimensionality reduction within each attention gate was achieved via (1 × 1 × 1) convolutions on both the guidance signal and the encoder feature map, which were then projected into a shared intermediate space, and combined through element-wise summation, followed by a ReLU activation. Subsequently, a (1 × 1 × 1) convolution and Sigmoid activation were used to generate an attention map and scale each spatial location to values between 0 and 1. The encoder feature map is then multiplied by the attention map and concatenated with the upsampled decoder feature map along the channel dimension.

#### 2.4.3. SegResNetVAE

The SegResNetVAE is a 3D Variational Autoencoder (VAE) built upon the SegResNet architecture introduced in [17]. The encoder has five down-sampling stages, wherein the spatial dimensions are reduced by a factor of two using strided convolutions. The number of residual blocks in each stage is defined as (1, 3, 3, 3, 4). Each residual block comprises instance normalization, LeakyReLU activation, and two (3 × 3 × 3) convolution layers. The initial number of filters is set to 64, which doubles at each downsampling stage and reaches 1024 in the bottleneck. The VAE branch estimates both the mean (*µ*) and standard deviation (σ) of a latent representation with a dimension of 512, which helps with regularization. The decoder uses transposed convolutions to reconstruct spatial dimensions, incorporating skip connections from the encoder to maintain spatial context. The upsampling path contains (3, 3, 3, 1) residual blocks at each stage. Instance normalization, LeakyReLU activation, and a final (1 × 1 × 1) convolution layer are employed to produce the output segmentation map.

#### 2.4.4. Swin UNETR

The Swin UNETR architecture [18] employs a hierarchical Swin Transformer with shifted windows as encoder and a CNN-based decoder. The feature representations at different stages of the encoder are passed to the corresponding level in the decoder using the skip connections.

Our implementation uses a patch size of (2 × 2 × 2). A linear embedding layer projects the raw features of each patch to an embedding space of size (*C* = 72). The encoder has four stages. Each stage comprises a patch merging block to reduce the spatial dimensions of the feature map by a factor of two, along with two Swin Transformer blocks. A Swin Transformer block incorporates a shifted window-based multi-head self-attention (WMSA) module, a 2-layer Multilayer Perceptron (MLP), and a Gaussian Error Linear Unit (GELU) nonlinearity in between. The initial number of attention heads is set to 6 and doubles at each stage.

As mentioned, Swin UNETR features a CNN-based decoder. At each stage of the decoder, the output features are reshaped and processed through a residual block. Each residual block comprises two (3 × 3 × 3) convolutional layers with instance normalization. Upsampling of feature maps is achieved through a deconvolutional layer, which is then concatenated with the output of the previous stage and processed by another residual block. Finally, a (1 × 1 × 1) convolutional layer is implemented to generate the segmentation map.

### 2.5. Implementation details and hyperparameters

All models were trained to predict six tissue classes: blood vessels, cells, vacuoles, axons, boundary class, and background. We used the boundary class around the axons to address under-segmentation, known as the “merging axons” problem, where adjacent axons are mistakenly identified as a single instance. The boundary class was generated by dilating the axonal masks with a 1 × 1 × 1 structuring element and then subtracting the original axon masks from the dilated mask. Due to memory limitations, the input data to the models was 3D patches rather than entire XNH volumes. Since XNH scans have a cylindrical FOV, the image matrix includes non-tissue regions. To address this, a tissue mask is generated to exclude these regions. 3D patches were sampled randomly from this cylindrical FOV using the tissue mask. This process was facilitated through the TorchIO library [21].

Patch size compromises the 3D size of detected anatomical features and GPU memory, even when using high-memory GPUs such as the NVIDIA A100. To investigate the effect of patch size on the segmentation performance of the model, we first tested three different patch sizes: 64 × 64 × 64 (PS64), 128 × 128 × 128 (PS128), and 140 × 140 × 140 (PS140) using our U-Net model. PS140 was the maximum that the GPU could accommodate. Finally, as a compromise between memory and performance, we chose PS128 to compare different model architectures and loss functions.

Before applying the XNH data to the models, all image volumes were intensity-normalized by their means and standard deviations. The following settings were used during the training process for all models: Adam optimizer, an initial learning rate (LR) of 0.00005–0.0001 decayed through a polynomial LR scheduler (power = 0.9), a batch size of 4–5, and 800 epochs after which early stopping was applied. Data augmentation was performed on-the-fly to prevent overfitting and increase data diversity during model training. The augmentation pipeline combined random flipping with a 50% probability and one of two affine transformations: (1) a zoom-out transformation with scaling factors in the range [0.3, 1.0] and random rotations between -180° and 180° (80% probability); or (2) a zoom-in transformation with scaling factors in the range (1.0, 3.0] and random rotations between -180° and 180° (20% probability). All augmentations were performed in 3D across all spatial axes. The training was conducted on an NVIDIA A100 cluster with 80GB of memory, provided by the High-Performance Computing (HPC) services at the Technical University of Denmark (DTU).

### 2.6. Segmentation loss functions

We explored three different loss functions for the segmentation task, including cross entropy (*L*_CE_) as a distribution-based loss, generalized dice loss (*L*_GDL_) as a region-based loss, and generalized dice focal loss (*L*_GDFL_), which is the weighted sum of *L*_GDL_ and ‘focal loss (*L*_FL_), with equal contribution (*ε* = 0.5). The loss functions are expressed as follows.

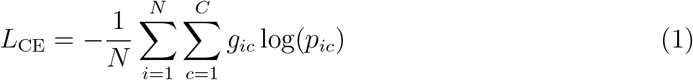

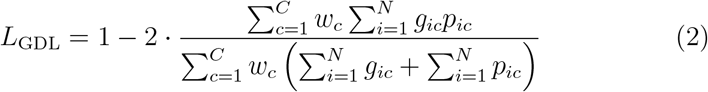

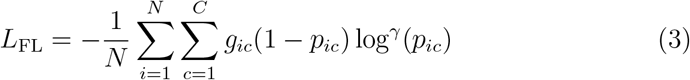

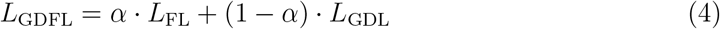

where *N* and *C* denote the number of pixels and classes, respectively. *p*_*ic*_ denotes the predicted probability of voxel *i* belonging to class *C*, and *g*_*ic*_ is the corresponding GT. *w*_*c*_ is the class weight, inversely proportional to the squared sum of GT voxels in class *C*. γ is the focusing parameter (γ ∈ [1, 3]) [22], which helps to emphasize more on hard-to-classify regions, with the default value of γ = 2 [22] implemented in this work.

#### 2.6.1. Deep supervision

Our U-Net and Attention U-Net architectures also incorporate deep supervision and auxiliary losses for better gradient flow [22]. To implement this, we downsampled the GT labels using nearest-neighbor interpolation to match the dimensions of the outputs extracted from different depths of the decoder. We then weighted these auxiliary losses based on their corresponding decoder level. The final loss function was computed as a weighted sum of losses at different depths of the decoder part, as follows.

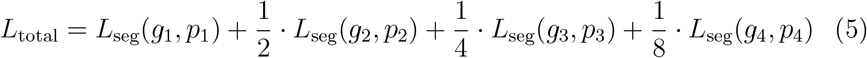

where *g*_*i*_ and *p*_*i*_ represent the resized GT and predictions at decoder level *i*, respectively. *L*_seg_ denotes the segmentation loss (*L*_CE_, *L*_GDL_, *L*_GDFL_) as introduced in the previous section.

#### 2.6.2. VAE loss for SegResNetVAE

For SegResNetVAE, the loss function includes three terms: *L*_seg_, the segmentation loss; *L*_MSE_, the *L*_2_ loss on the VAE branch output (*I*_pred_) to match the input image (*I*_input_); and *L*_KL_, the VAE penalty term as [17], a Kullback–Leibler (KL) divergence between the estimated normal distribution 𝒩 (*µ, σ*^2^) and a prior distribution 𝒩 (0, 1):

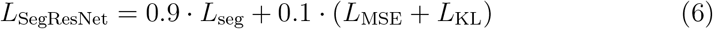

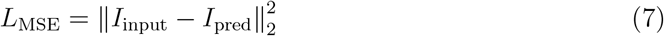

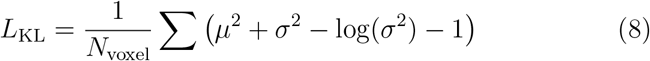

### 2.7. Performance evaluation

To measure the evaluation metrics, we compared the model prediction on the test dataset with EGT. We reported overall accuracy, recall, precision, Dice score, intersection over union (IoU) to evaluate the performance of all trained models following the equations below. TP, TN, FP, and FN represent true positives, true negatives, false positives, and false negatives, respectively.

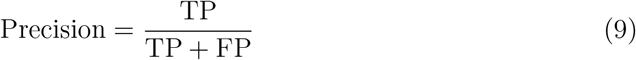

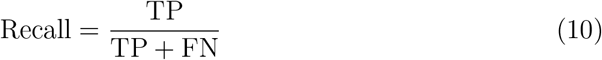

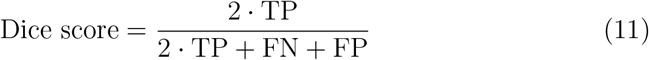

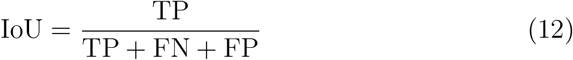

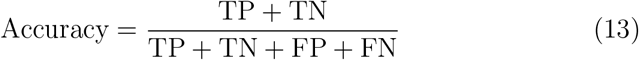

### 2.8. Quantification

While morphological quantification is beyond the scope of this work, we conducted an initial analysis to assess how microstructural feature sizes are extended by incorporating the expert-in-the-loop strategy. Analysis was conducted on GT and the final EGT. For cells and vacuoles, we first performed a connected component analysis for instance segmentation and then calculated the volume of each component based on the number of voxels it occupied. Components smaller than 30 µm^3^ in volume were excluded from further analysis.

For axons, we first performed a connective component analysis to separate individual instances. Next, we excluded those axonal segments spanning less than 100 voxels and shorter than 10 µm in length. For the rest, we extracted the centerlines through skeletonization. We measured the diameter along the centerline at regular intervals of 500 nm in the cross-sections perpendicular to it. An ellipse was fitted to each of these cross-sections, and the equivalent diameter of the fitted ellipse was then calculated as a measure of the axonal diameter [4]. These locally calculated diameters were averaged to obtain the mean axonal diameter. The curve length of the centerline is also used to define the axonal length. For blood vessels, we calculated the mean diameter of each branch using the same method described above to calculate the mean axonal diameter. All these calculations were performed in MATLAB R2023a and Python 3.10.10.

## 3. Results

### 3.1. Impact of EGT on the model performance

In the initial training, using only the sparse GT labels, we achieved Dice scores of 0.81, 0.68, 0.65, and 0.48 for blood vessels, vacuoles, cells, and axons, respectively. Upon expanding the dataset and incorporating EGT labels, we observed clear improvements, with Dice scores increasing to 0.94 for blood vessels, 0.84 for vacuoles, 0.7 for cells, and 0.93 for axons. The overall accuracy was also improved from 0.94 to 0.975. The qualitative comparison of the segmentation performance of the model in the initial and final rounds of training, as shown in Fig. 2, confirms the quantitative gains observed in terms of Dice score. In particular, improved model performance in Fig. 2 could be seen by many more segmented smaller axons, better segmentation of larger axons, and fewer over-segmentation errors mainly related to intensity variations inside the axons (orange arrows in Fig. 2). A better delineation of blood vessels and vacuoles were also achieved, which were often misclassified by the initial model (red arrows in Fig. 2). The detection of cells also improved, with the final model identified some of small cells that were missed in the initial round of training with sparse GT labels (white arrows in Fig. 2). Figure 3 illustrates how the feature size for different microstructures is extended using expert-in-the-loop label refinement, particularly for the axons. In the initial GT, we had annotations for a few axons with a diameter ranging from ∼2 to 4 *µ*m and a maximum length of ∼170 *µ*m. However, EGT includes many more axons with diameters between ∼1 and 8 *µ*m and lengths up to 600 *µ*m. We must acknowledge that, even though the model was able to segment some of the smaller axons (*<*1 *µ*m), the majority were left out during the post-processing and cleaning stage, as they were too small to be reliably quantified at the given resolution of 375 nm. According to the histograms presented in Fig. 3, it is evident that EGT also captures a broader range of sizes for cells and vacuoles. Notably, we observed larger clusters of cells and larger vacuoles in EGT, reaching volumes up to 22000 *µ*m^3^ and 1400 *µ*m^3^, respectively, which were not present in GT. Indeed, expert-in-the-loop refinement not only increased the number of detected instances but also enhanced the representativeness of the dataset by including more morphologically diverse examples that were previously underrepresented in the GT.

**Figure 2:**
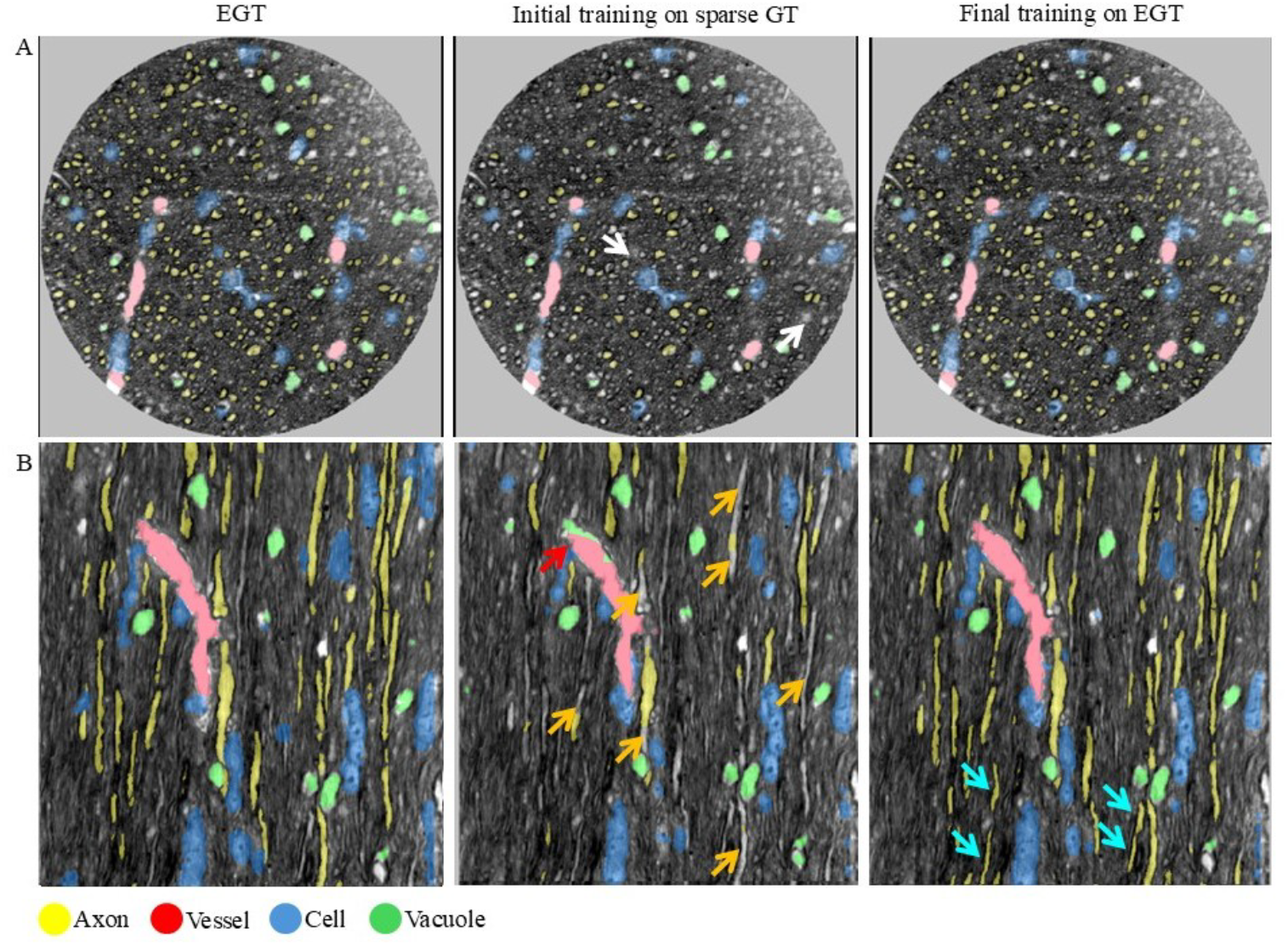
Qualitative comparison of the model performance in the initial and final rounds of training. From left to right: Extended ground truth (EGT), initial training on sparse GT labels, and final performance following expansion of the dataset and integration of EGT using expert-in-the-loop training. A and B show the selected axial (#1330) and sagittal (#291) slices from the test dataset in the corpus callosum (CC), respectively. While improvement in segmentation performance of the model was observed for all classes, the most notable enhancement was seen in axonal segmentation. The orange arrows indicate the poor performance of the initial model in axonal segmentation. The red arrow indicates the misclassification error in blood vessel segmentation in the initial training. Some of the small cells were also missed by the initial model (indicated by white arrows), which were detected by the final model trained on EGT. Many new axons are also segmented by the final model that were not annotated in EGT (cyan arrows). Results are shown for a 3D U-Net model trained with generalized dice focal loss (GDFL).

**Figure 3:**
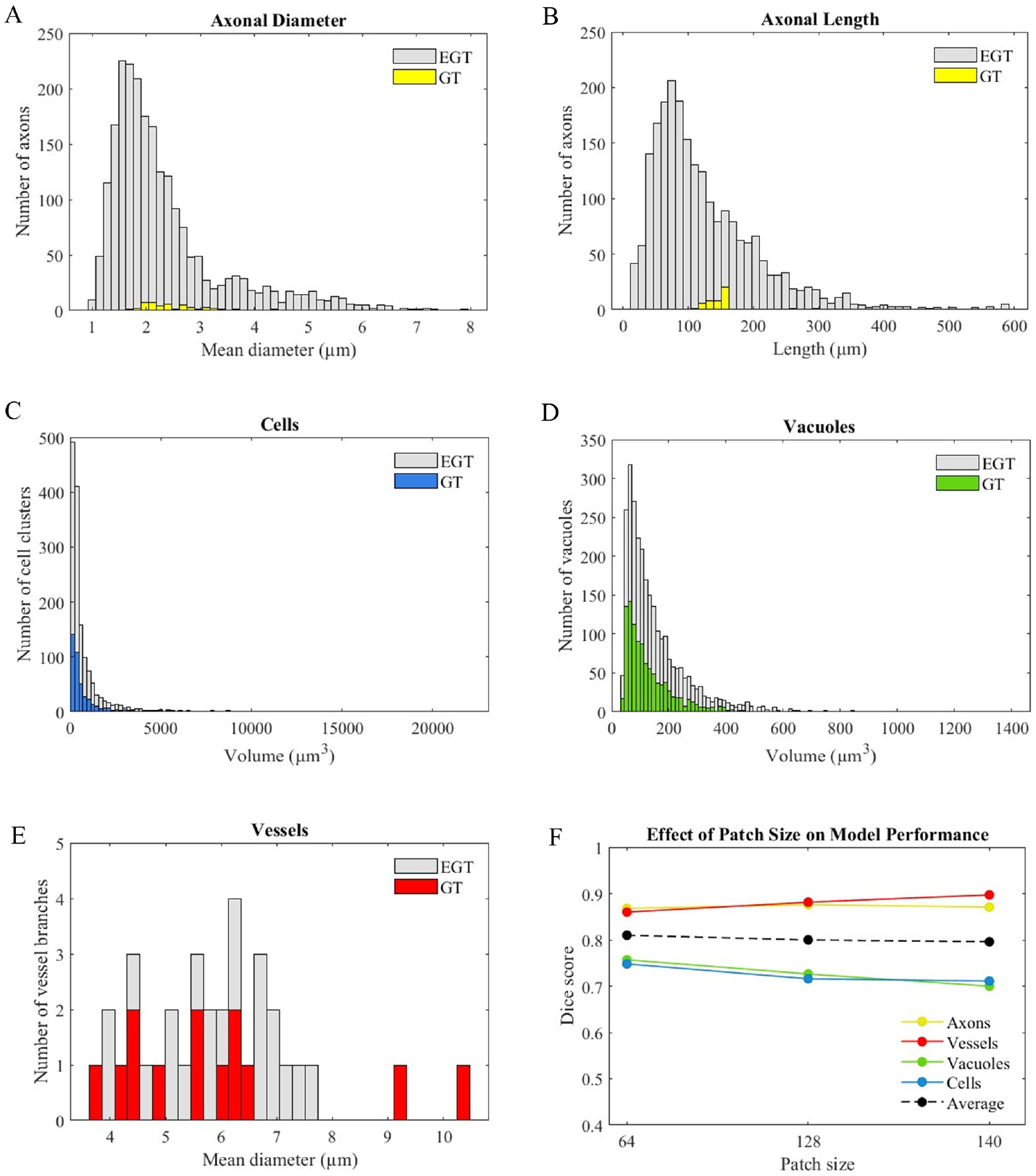
(A–E) Distributions of various structural features across different microstructures in initial ground truth (GT) and extended ground truth (EGT). The difference between EGT and GT histograms is more pronounced for the axons than for the other microstructures. (F) Class-wise Dice score evaluation across different patch sizes. Larger patch sizes surge the performance in the segmentation of larger microstructures such as vessels and axons. However, performance for vacuoles and cells declines when increasing the size of training patches. The average Dice scores across all classes of microstructures are indicated with the dashed black line.

### 3.2. Patch size

We explored the effect of patch size on the segmentation performance of the model across different microstructures. To achieve this, we trained our 3D U-Net model using three different patch sizes: 64 × 64 × 64 (PS64), 128 × 128 × 128 (PS128), and 140 × 140 × 140 (PS140). For this experiment, training was only performed on the CC dataset, and GDFL was used as the loss function. As shown in Fig. 3F, the average Dice scores were 0.81 for PS64, 0.80 for PS128, and 0.796 for PS140, respectively. Using smaller patches (PS64), the model performed well on small structures such as cells and vacuoles while struggling to capture larger components, such as blood vessels and larger axons in CS. A qualitative comparison of model performance on the test dataset from the CS region for different patch sizes is illustrated in Fig. 4. As shown in Fig. 4A, the models trained on bigger patches of PS128 and PS140 failed to detect some of the small cells (white arrows). However, these were identified when employing smaller patches of PS64.

**Figure 4:**
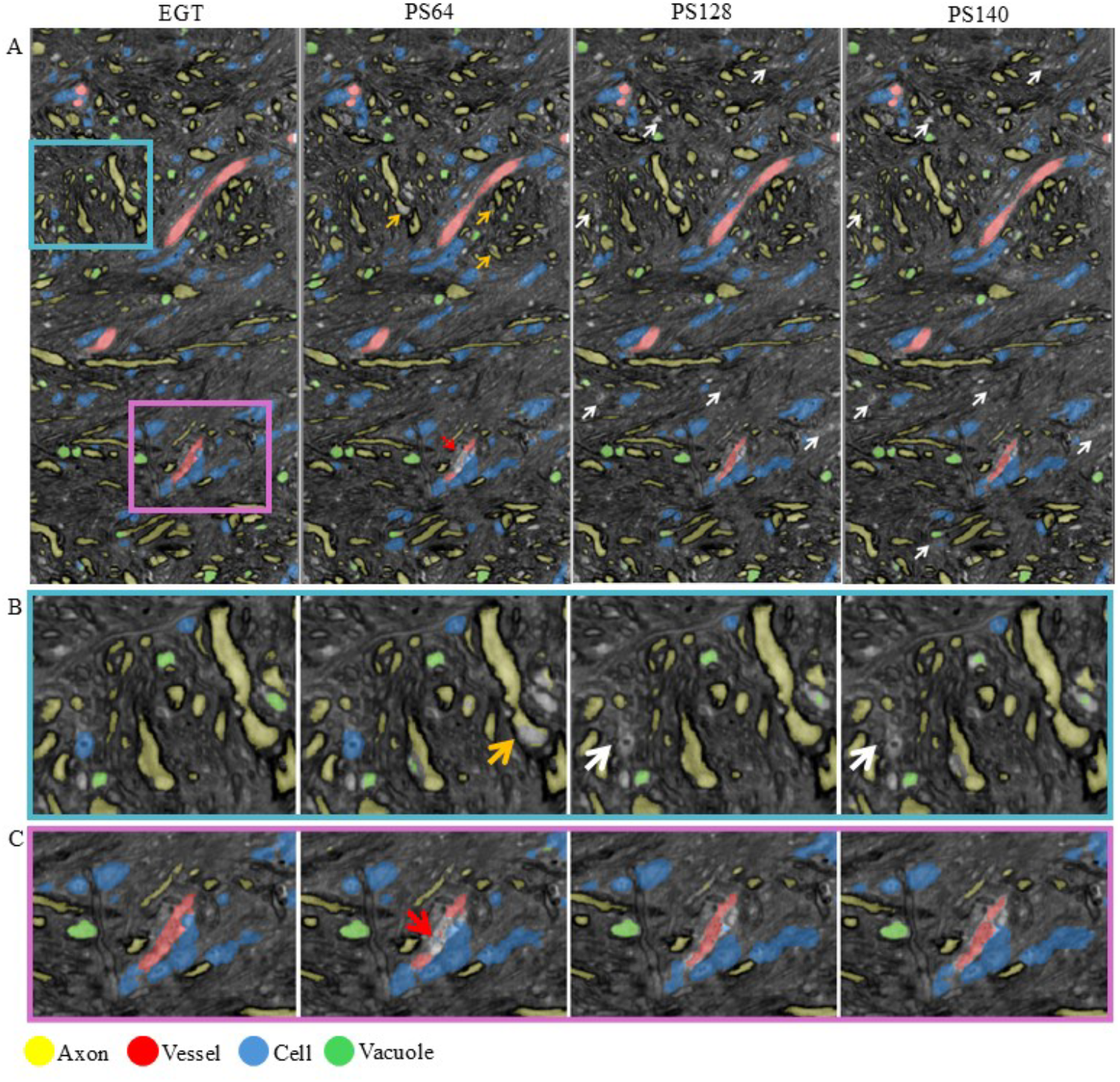
Influence of patch size on the segmentation performance of the model. Results are shown for a sagittal slice (#237) from the centrum semiovale (CS), along with zoomed-in views of selected regions (B and C). The small cells that were missed by the models trained on larger patch sizes of PS128 and PS140 are shown by white arrows, whereas the model trained on smaller patches of PS64 detected these instances. Red and orange arrows point out the poor segmentation of blood vessels and some of the large axons when using small patches of PS64 for training.

In contrast, increasing the patch size led to better segmentation of blood vessels and larger axons, particularly in CS, by providing the model with more contextual information. This improvement, compared to PS64, is highlighted by orange and red arrows in Fig. 4B and C. However, it does so at the cost of greater computational requirements and lower segmentation performance in segmenting smaller microstructures. Increasing the patch size beyond PS128 did not offer additional benefits. Consequently, a compromise was made between computational feasibility and segmentation performance across all microstructural classes, and PS128 was selected for the rest of our experiments.

### 3.3. Model architectures and loss functions

Using EGT, we studied a set of different architectures and loss functions. Each model was trained by three different loss functions: CE, GDFL, and GDL. The segmentation performance of all trained networks was evaluated on 3D annotations from unseen test datasets in CC and CS regions and presented in Fig. 5. Dice scores for different classes are also summarized in Table 1. Overall accuracy, class-wise recall, precision, and IoU for all networks are also provided in the supplementary information (SI). Only slight differences in performance were observed when training the same model with different loss functions, as shown in Table 1. Note that the average Dice scores in Table 1 are only computed for microstructures and do not include background values. The overall performance of U-Net in terms of Dice score was 0.849, 0.85, and 0.856 for CE, GDFL, and GDL, respectively. Attention U-Net showed the least dependency on the loss function with Dice scores of 0.844, 0.848, and 0.843 for CE, GDFL, and GDL, respectively. The Dice scores of SegResNetVAE were calculated to 0.831, 0.85, and 0.843 when training with CE, GDFL, and GDL, respectively. Swin UNETR, the transformer-based architecture, achieved a Dice score of 0.807 for CE, 0.831 for GDFL, and 0.818 for GDL. Results presented in Table 1 have shown that among all tested networks, U-Net and SegResNetVAE achieved the highest Dice scores. While better precision was achieved for SegResNetVAE compared to U-Net (regardless of loss function), average training was faster for U-Net compared to SegResNetVAE. Figure 6 shows the final segmentation results of the U-Net trained with GDFL for two different WM regions, following expert proofreading and cleaning.

**Table 1:**
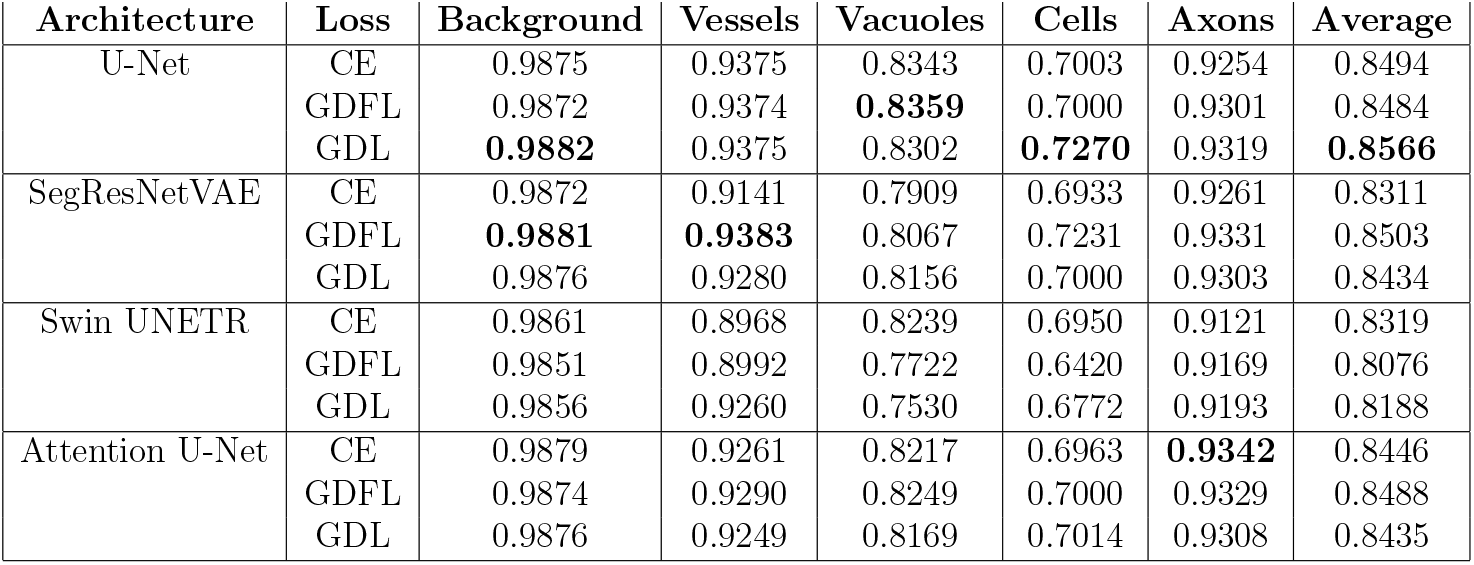
Comparison of Dice scores across different architectures and loss functions. All models were trained in 3D using an isotropic patch size of 128 × 128 × 128. The average score computed over microstructures, excluding background. Values in bold show the best results achieved for each class. Abbreviations: GDL: generalized dice loss, CE: cross entropy, GDFL: generalized dice focal loss.

| Architecture | Loss | Background | Vessels | Vacuoles | Cells | Axons | Average |
| --- | --- | --- | --- | --- | --- | --- | --- |
| U-Net | CE | 0.9875 | 0.9375 | 0.8343 | 0.7003 | 0.9254 | 0.8494 |
|  | GDFL | 0.9872 | 0.9374 | **0.8359** | 0.7000 | 0.9301 | 0.8484 |
|  | GDL | **0.9882** | 0.9375 | 0.8302 | **0.7270** | 0.9319 | **0.8566** |
| SegResNetVAE | CE | 0.9872 | 0.9141 | 0.7909 | 0.6933 | 0.9261 | 0.8311 |
|  | GDFL | **0.9881** | **0.9383** | 0.8067 | 0.7231 | 0.9331 | 0.8503 |
|  | GDL | 0.9876 | 0.9280 | 0.8156 | 0.7000 | 0.9303 | 0.8434 |
| Swin UNETR | CE | 0.9861 | 0.8968 | 0.8239 | 0.6950 | 0.9121 | 0.8319 |
|  | GDFL | 0.9851 | 0.8992 | 0.7722 | 0.6420 | 0.9169 | 0.8076 |
|  | GDL | 0.9856 | 0.9260 | 0.7530 | 0.6772 | 0.9193 | 0.8188 |
| Attention U-Net | CE | 0.9879 | 0.9261 | 0.8217 | 0.6963 | **0.9342** | 0.8446 |
|  | GDFL | 0.9874 | 0.9290 | 0.8249 | 0.7000 | 0.9329 | 0.8488 |
|  | GDL | 0.9876 | 0.9249 | 0.8169 | 0.7014 | 0.9308 | 0.8435 |

**Figure 5:**
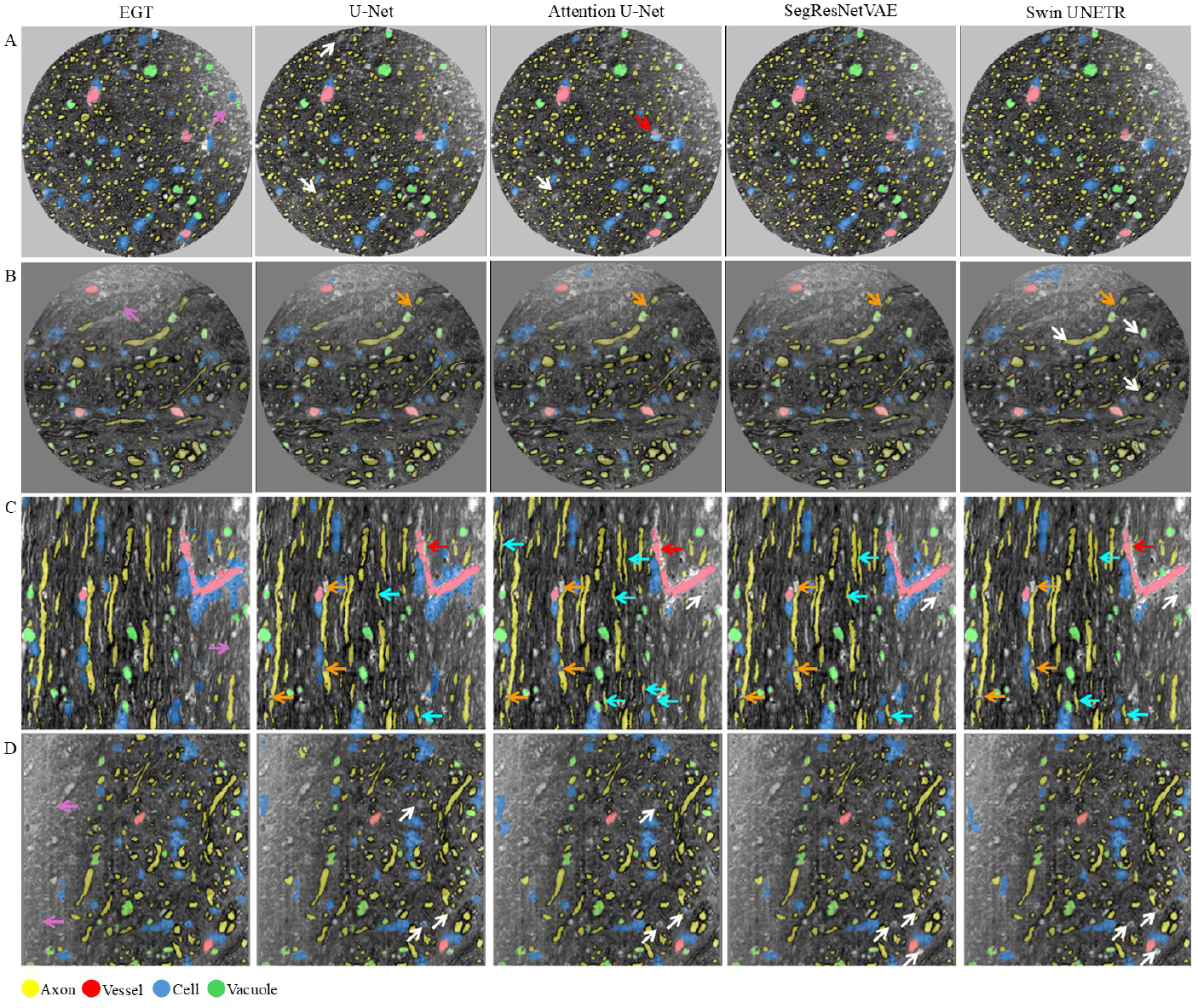
Segmentation performance of different model architectures on the test dataset from the corpus callosum (CC) and centrum semiovale (CS) regions. A and B illustrate the selected axial views from the CC (#1448) and CS (#1478), respectively. C and D display sagittal views from the CC (#215) and CS (#205), respectively. Red arrows point out the blood vessels misclassified by the networks. White arrows highlight incomplete and fragmented segmentations of cells and vacuoles. Some of the over-segmentation errors in axons are shown with the orange arrows. Pink arrows on EGT show the low-contrast regions affected by staining issues. The models also identified new instances of axons (cyan arrows) that were not previously annotated in the extended ground truth (EGT). Note that results are shown for models trained with generalized dice focal loss (GDFL) as the loss function.

**Figure 6:**
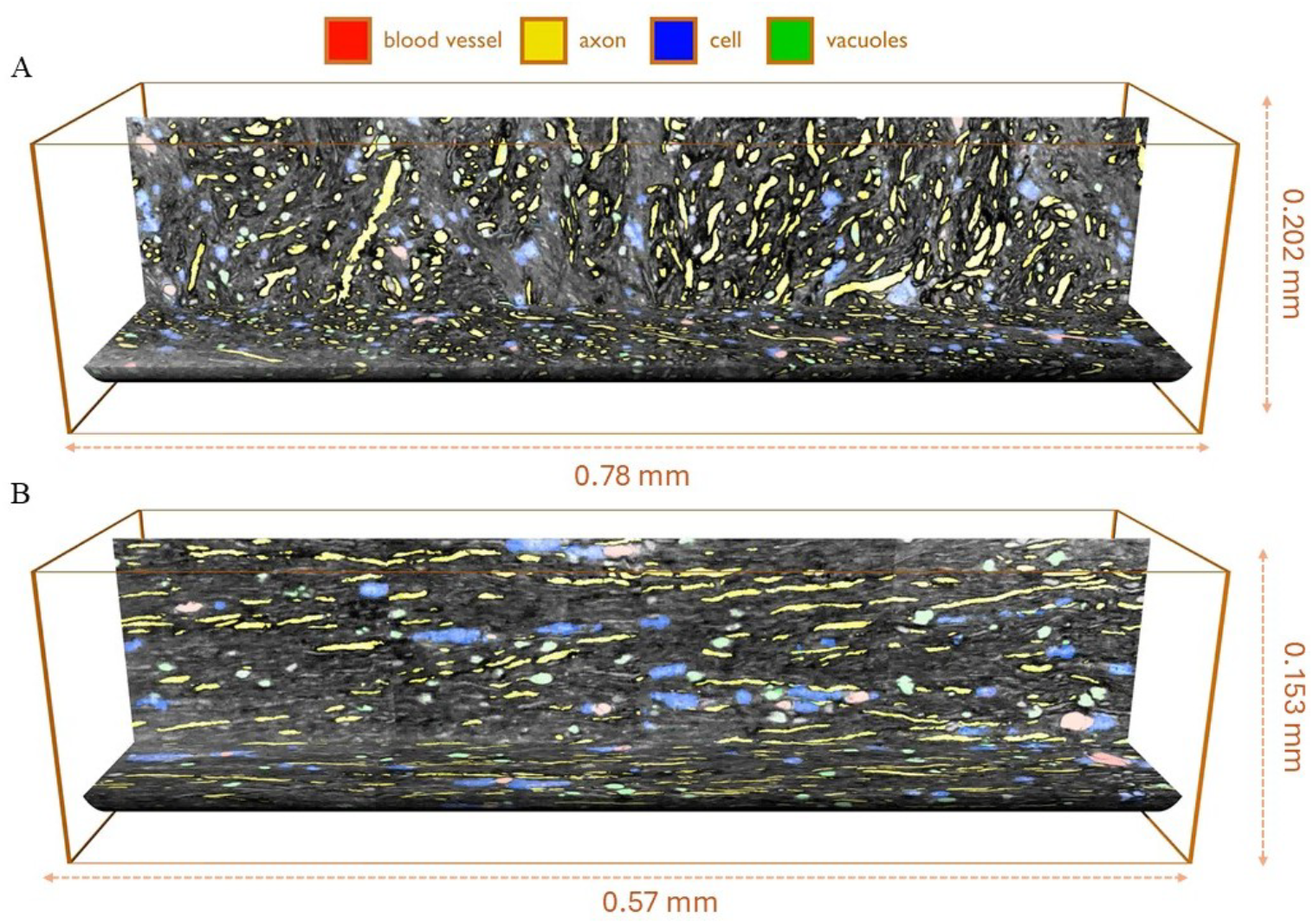
Final segmentation results for extended FOV from two different regions of the WM. A and B represent the segmented microstructures (axons, cells, vessels, and vacuoles) in centrum semiovale (CS) with relatively complex axonal morphology and corpus callosum (CC) with simple and parallel axonal orientation, respectively.

Figure 5 presents a qualitative comparison of different model architectures on the test dataset from both WM regions. As shown, all models achieved decent results in multi-class segmentation of WM microstructures. Some small axons were also detected by the models, especially by the Attention U-Net, despite not being annotated in the EGT (cyan arrows in Fig. 5). Upon visual inspection, a few segmentation errors were also observed, especially for the cells, which explains their relatively lower Dice scores across all models. We noticed that some small cells remained undetected by all models (indicated by white arrows), particularly those located at the corners of the FOV or within poorly stained areas, likely due to insufficient contextual information in these regions. Additional segmentation errors in axons (highlighted by orange arrows in Fig. 5) were also observed. These errors were more evident close to other microstructures, especially vacuoles, where axonal diameters typically narrow. In these scenarios, the segmentation performance declined for all models.

## 4. Discussion

Combining a DL-based approach with expert-in-the-loop label refinement, we achieved 3D full segmentation of four different types of microstructures with an average Dice score of 0.856 in large FOV XNH volumes up to 202 × 202 × 780 µm^3^ of primate brain WM samples. Starting with a sparse subset of 3D annotations of 54 axons within a “simple” tissue region, we segmented over 2300 instances of long-range axons in 3D. These include both straight axons from the corpus callosum (CC) and axons from the crossing fiber region (CS) with complex and varying morphological features (e.g., diameter, length, and directionality). Our method achieved a Dice score of 0.931 in 3D axonal segmentations, comparable to the results reported in 2D [7, 10, 8] and 3D EM studies [13, 12]. However, unlike most previous works that focused only on axonal segmentation, we advanced the scope to 3D multi-class segmentation of various microstructures in the WM. In virtue of XNH wide FOV, we were not only able to study long-range axons up to 600 µm but also to segment and analyze additional microstructures. We segmented the whole blood vessels, thousands of vacuoles, and cells from both WM regions. This work represents a significant step toward comprehensive 3D analysis of WM microstructures using the wide FOV XNH technique, addressing the gap left by axon-centric studies while offering a more holistic view of WM microarchitecture and organization. As we extended the labeled dataset, we sought to train and benchmark different architectures and loss functions to identify the most effective model in this segmentation context. This model serves as the foundation for future large-scale segmentation efforts as additional samples become available and can be readily shared with the broader community to support research on similar datasets.

### 4.1. The expert-in-the-loop strategy effciently extends the 3D feature space

Applying the expert-in-the-loop label refinement resulted in a 23% improvement in the overall segmentation performance of the model in terms of Dice score compared to training on the sparse initial GT. While this improvement is not surprising, as more labels were introduced, the key advantage of expert-in-the-loop lies in streamlining the annotation process. This is particularly important in large-scale annotation tasks. This way, we increased the labeled dataset by sixfold within just a few iterations, where each step required a lesser degree of expert annotation and editing. The expert-in-the-loop strategy has also proven effective in EM [13, 23] and biomedical segmentation [24, 25], speeding up expert annotation time and improving segmentation robustness, which aligns well with our observations.

We also observed that the level of improvement with the expert-in-the-loop strategy varies among classes. The gain was more pronounced for axons (∼50%) than for other microstructures (∼10%). It was expected, as axons had a worse starting point than the other microstructures. Therefore, we achieved a relatively lower Dice score for the axons in the first round of training. Moreover, the initial GT labels for the axons only captured a narrower range of morphological variations. Although we applied data augmentation (i.e., scaling) to artificially extend the size range of anatomical features during training, the expert-in-the-loop approach still improved the segmentation performance. Indeed, the real tissue morphology for the axons was out-of-distribution and poorly represented in the initial GT. In the initial training, we only had annotations for the 54 largest axons from CC, which were predominantly parallel, straight, and smaller-diameter axons. In contrast, the second dataset, from the CS region, features a more diverse axonal morphology and organizational complexity. CS is mainly occupied by non-straight, crossing, and tortuous axons, including the so-called giant axons with a diameter of up to 8 µm. These CS axons, being out-of-distribution in initial training, were poorly handled by the initial model trained solely on a small volume of the CC. Therefore, by expanding the training dataset and including annotations from the CS using the expert-in-the-loop strategy, we improved the segmentation performance, particularly for those axons with more complex geometries. It also helped to reduce the over-segmentation errors caused by intra-axonal intensity variations, which particularly exist in CS. Compared to the axons, initial GT labels for blood vessels, cells, and vacuoles were sufficient to represent their typical morphology. As the morphology and size of these microstructures are more consistent across different regions of WM (unlike axons), indicated by their histograms in Fig. 3, extending their annotations yielded only moderate performance gains. For example, cells and vacuoles are mostly spherical and fall within a narrow size range of approximately ∼5–10 µm. Blood vessels are also consistent in size and shape within a specific segment or branch. In contrast, axonal diameters can vary significantly from 0.1 µm to 8 µm [4, 26].

### 4.2. No specific model architecture outperformed the others

The expert-in-the-loop strategy was applied exclusively to a 3D U-Net architecture. It was a deliberate choice, as U-Net has a simple architecture and is therefore more robust when trained on a sparsely labeled dataset. With the enhanced dataset in place, we aimed to train the most effective and generalizable model possible. Therefore, we explored a set of alternative architectures, including Attention U-Net, SegResNetVAE, and Swin UNETR. By benchmarking various architectures, we aimed to identify the most performant model for our data and provide a valuable point of comparison for others developing models on a similar segmentation task.

To make a fair comparison among different architectures, we adjusted the number of parameters in each network to be comparable (except for the transformer). The trainable parameters were set to be ∼90M, 91M, 94M, and 163M for U-Net, Attention U-Net, SegResNetVAE, and Swin UN-ETR, respectively. While we found comparable performance among different model architectures, CNN-based models performed slightly better than the transformer. It is well established that transformer-based architectures typically require large datasets or large-scale pretraining to achieve comparable or superior performance to CNNs, as they lack the strong inductive bias of CNNs [27]. Transformers are also more memory-intensive and computationally expensive due to their self-attention mechanism, which limits their application, particularly for large-scale 3D segmentation tasks like ours. Moreover, we used the same settings (except for the learning rate) to train all models to minimize the impact of other parameters that could confound the results. More comprehensive fine-tuning of hyperparameters for each architecture may improve their performance further and better reflect their optimal capabilities.

We noted improved precision for almost all classes by incorporating attention gates. However, sensitivity was lower, leading to lower Dice scores. Improvement in precision was also observed for axons and cells when using SegResNetVAE. The segmentation performance of SegResNetVAE, as measured by the Dice score, was comparable to that of U-Net; however, incorporating a VAE branch adds complexity to the network architecture and increases training time.

No further improvements were achieved when using Swin UNETR. Similar findings have been reported in the Brain Tumor Segmentation (BTS), where the author evaluated different U-Net variants on a large training dataset and showed that a plain U-Net achieved slightly better Dice scores than the transformer and also performed similarly to SegResNetVAE, despite its simpler architecture [28].

Indeed, the marginal difference among models suggests that all models have sufficient capacity to learn the feature variation among all classes of microstructures. Moreover, all models demonstrated reliable segmentation performance as reflected in their Dice scores in Table 1. This is because the training dataset and EGT suffciently represent the statistical distribution of key feature variations such as size, intensity, and morphology across different classes within the types of samples included here. Extending training data beyond the CS and CC samples is unlikely to enhance the segmentation performance further, unless the feature information in the dataset improves. This could be achieved, for example, through better image quality or improved image resolution. Together, these factors determine the structural resolution limit. The structural resolution limit defines the 3D image matrix size, which must be suffciently fine-gridded to spatially sample the geometric features of interest for segmentation. The limit can be reduced by interpolation, which has a similar effect to Gaussian image blurring. Herein, we downsampled the original data to enhance the signal-to-noise ratio (SNR). However, in most EM studies, images were downsampled to match the resolution limit to relevant feature information [29] or to address the memory demands [12]. Hence, better SNR in our XNH data would increase our resolution limit without further downsampling or Gaussian filtering. However, patch size is finite and memory-dependent. Therefore, increasing the resolution limit without increasing patch size may come with the risk of losing broader contextual information, which can negatively impact the segmentation performance of the model as discussed in the following section on patch sizes.

We also trained each model architecture on EGT using three different loss functions. A minor difference was observed when training the same model architecture using different loss functions. Overall, region-based losses, such as GDL and GDFL, performed better than the distribution-based CE loss, but the improvements were marginal. In general, Dice-based losses are shown to be more robust across different segmentation tasks [30], particularly in tasks with highly imbalanced labels, as they rely on maximizing the overlap between predicted and GT labels rather than voxel-wise similarity. Another critical factor that can impact the class imbalance is the patch size and patch sampling strategy [30]. Herein, we employed a tissue mask to oversample the patches from the foreground region where labels were available. Moreover, the patch size was big enough to include labels from all classes. Together, these likely contributed to mitigating global class imbalance by providing a more balanced representation of different classes within the training samples. As a result, the performance differences between the loss functions remained marginal.

### 4.3. Patch size determines the range of feature sizes being segmented

We found a moderate relationship between input patch size and segmentation performance across structural scales. In general, changing the patch size changes the effective receptive field of the model. Training with smaller patches (e.g., 64 × 64 × 64) provided more local information, enabling the model to better capture fine-grained features in the images. As a result, structures such as cells and vacuoles, being small and densely distributed, were effectively segmented. This configuration was also faster and more computationally efficient. However, the main limitation of using small patches is the small receptive field, which limits the ability of the model to capture broader contextual information. This information is necessary for segmenting larger structures spanning across multiple patches, such as larger blood vessels and bigger axons, in our case.

In contrast, larger patches (e.g., 128 × 128 × 128) broaden the effective receptive field and provide more global context information required for segmenting larger structures. Yet, segmentation performance for the smallest structures tended to decline when increasing the patch size from PS64 to PS128. Another important point to mention is that even though overall segmentation performance showed a slight improvement when increasing the patch sizes (Fig. 2), this came with a significant increase in memory consumption and training time, approximately four times in our case. We should also note that our ability to experiment with even larger 3D patch sizes (e.g., 256 × 256 × 256) was limited by the available memory on the A100 graphics cards we used. Moreover, increasing the isotropic patch size from PS128 to PS140 offered no further benefit in our case.

Previous studies have also employed patch-based training rather than using whole 2D slices [11, 10] or entire 3D volumes [12, 13]. However, they often do not explicitly consider the importance of patch size, as most of these studies were performed on only axonal segmentation, which can also be captured effectively using small patches and smaller FOVs. However, our experiments highlight that selecting an appropriate patch size is a critical design decision. This is particularly important in multi-class segmentation where microstructural features span a wide range of spatial scales [5], as in our dataset.

### 4.4. Over- and under-segmentation: a limitation of image resolution

We achieved an excellent Dice score of > 0.9 in axonal segmentation. These results were consistent with visual inspection. Axonal boundaries are often delineated clearly by surrounding dark myelin layers, making them easier to segment even after a five-fold downsampling. However, over-segmentation or “split axons” is inevitable in axonal segmentation due to several reasons related to both image quality (e.g., staining and resolution limitation) and anatomical factors (e.g., tension caused by adjacent microstructures and also Nodes of Ranvier) [4]. For example, we observed that axonal diameters locally shrank to a few pixels in proximity to other microstructures like vacuoles or cells, as shown in Fig. 5. In such cases, the axon diameter falls below the resolution limit and appears fragmented. These segments will likely be removed during post-processing if they are too small. A similar pattern was also observed at the Nodes of Ranvier, where the axon narrows and there is no myelin around the axons, hence no boundary contrast, which makes the boundary detection difficult and splitting more likely [12]. These nodes of Ranvier, spaced roughly every 300 µm, define the length of the myelin sheath [4]. Ideally, an axon would be segmented as a single object, but segmentation breaks at the Nodes of Ranvier are expected and are a common issue in axonal segmentation [12]. Over-segmentation can be tackled by implementing further post-processing steps or expert proofreading. For instance, Plebani et al. reported increased F1-scores by employing expert evaluation following U-Net segmentation of peripheral axons while saving up to 80% of segmentation time [11]. In our study, we also implemented expert proofreading to correct ambiguous boundaries, Nodes of Ranvier, and other possible errors caused by staining and image quality. Interestingly, despite the lower resolution in this study, we could segment very long axons, owing to the wide FOV provided by XNH. We segmented long-range axons with length up to 600 µm, several times longer than those reported in higher resolution large FOV EM datasets (20, 100, and 200 µm) [29, 12, 13].

Another frequently reported issue in similar studies is under-segmentation or “merging axons”, where adjacent axons appear as a single object due to segmentation errors. We mitigated this by defining a boundary class around axons using morphological operations to highlight the surrounding myelin rim. The same strategy has been used in previous 3D axon [11] and cell nucleus [12, 15] segmentation studies.

Similar to axons, a reliable performance was achieved in segmenting blood vessels and vacuoles, as they also appear as high-intensity structures with well-defined boundaries. However, the cells had lower Dice scores than the other microstructures. All models yielded similar segmentation patterns as in Table 1. This discrepancy between EGT and model prediction could be attributed to the blurred and indistinct cell boundaries, which are challenging to define at the given resolution even for the expert raters. Cell membranes are lipid-based and extremely thin (∼10 nm) [31], therefore difficult to resolve even at 50 nm EM resolution [12]. In our downsampled data, these boundaries are even more obscured, and adjacent stained tissue becomes a proxy for the cell edge. As a result, all models struggle in areas where contextual information is limited, particularly near the edges of the FOV. Notably, boundaries were more blurred in the CS than in the CC sample due to the lower native resolution of 100 nm versus 75 nm, and thus, there were differences between expert defined boundries and model prediction. However, visual inspections showed good agreement between model output and anatomy. Similar challenges regarding cell segmentation have been reported in previous studies. Tong et al. [32] found that CNN-segmented cells were smaller but more consistent than those annotated by humans. Moreover, based on their analysis, cell boundaries defined by human annotators exhibited significant variability in repeated annotations, while the results of CNN segmentation were highly consistent.

Overall, our findings emphasize that the resolution limit is critical in instance segmentation, as it affects how well object boundaries can be identified and annotated for training. Moreover, the image resolution limit should not be confused with the image resolution itself. Image resolution is an absolute measure of voxel size. Resolution limit is relative and depends on how well the structures of interest can be sampled and separated within the image. Large structures can be accurately resolved even at lower image resolutions; small structures cannot. Since similar segmentation performance was found across all models, it suggests that all architectures are learning comparable spatial features, much like human annotators, whose performance also depends on boundary clarity. In general, the models appear to learn spatial shape information that helps distinguish different classes of microstructures with similar intensity profiles—for example, tube-like axons versus spherical-like cells. As a result, segmentation performance drops if anatomical boundaries or shape information cannot be robustly resolved due to the image resolution limit, especially for structures such as cells or small axons. Therefore, in future applications of DL for segmentation, careful consideration of the resolution limit in relation to the anatomical features of interest is essential for both model training and evaluation.

### 4.5. Limitations and future perspectives

We extended the GT labels based on very sparse initial annotations, trained a set of models, and demonstrated their comparable segmentation performance, but only on two samples (CC and CS) obtained from the WM of a monkey brain. We believe that the microstructural features the models learn are generalizable across the whole WM. Modest improvements in cell, vacuole, and blood vessel segmentation performance in Dice scores were observed when expanding from a single sample to both samples. Indeed, the axon class showed a clear improvement when including the CS sample, as it captures a larger part of the axon size distribution along with morphological variations known to exist in the brain [33]. Hence, the models are expected to generalize across the entire WM, which our preliminary segmentation results in other brain regions support (in preparation).

Generalizability depends on image quality. Both samples were stained individually, and visual inspection revealed similar image intensity and contrast; we performed intensity normalization across the two samples to minimize such variations. However, we excluded poorly stained regions from our evaluation [5], as segmentation performance dropped drastically in those areas. Recently, we have improved the sample preparation process, specifically minimizing surface scratches on the EPON samples, which significantly increases image quality toward EM standards. With these improvements, the five-times downsampling applied here to boost SNR would no longer be necessary, and therefore, microstructural boundaries appear clearer at the original image resolution. This improvement shifts the resolution limit downward and aids in segmenting smaller axon diameters. The price of better image quality—i.e., higher image resolution (no downsampling)—is the need for larger patch sizes to cover the full range of anatomical structures, thus requiring more memory, which we already faced as a limitation. The common solution is downsampling, as used in our presented configurations, or the adoption of new hierarchical network architectures capable of handling multiple levels of image resolution and patch sizes, each optimized for a particular range of structural geometries, such as HookNet [34] or cascade multi-resolution models [35]. Moreover, our samples are derived from a healthy monkey brain; it would be interesting to see how well the model generalizes to samples from other species or diseased brain tissues.

## Acknowledgements

This research was supported by funding from the European Research Council (ERC) under the Horizon Europe research and innovation program, provided through grant agreement No. 101044180 (Principal Investigator: T.B.D.). We acknowledge the ESRF for providing beamtime for experiment LS2702 at ID16A, from which parts have been made publicly available ([4, 5]) and utilized in this work. We also thank Thomson T.G. for help with data visualization.

## Author Contributions

M.A. and T.B.D. designed the study and interpreted the results. M.A. implemented the methods, performed the experiments, and analyzed the data. M.A. wrote the manuscript. T.B.D. and H.M.K. revised the manuscript. H.M.K. and M.A. contributed to the visualizations and figures. H.M.K. and M.L.B.T. contributed to the methodologies. All authors read and approved the final manuscript.

